# The microbiome mediates subchondral bone loss and metabolomic changes after acute joint trauma

**DOI:** 10.1101/2020.05.08.084822

**Authors:** Alyssa K. Hahn, Cameron W. Wallace, Hope D. Welhaven, Ellen Brooks, Mark McAlpine, Blaine A. Christiansen, Seth T. Walk, Ronald K. June

## Abstract

**Objective:** To compare the early responses to joint injury in conventional and germ-free mice.

**Design:** Post traumatic osteoarthritis PTOA was induced using a non-invasive anterior cruciate ligament rupture model in 20-week old germ-free (GF) and conventional C57BL/6 mice. Injury was induced in the left knees of n=8 GF and n=10 conventional mice. To examine the effects of injury, n=5 GF and n=9 conventional control mice were used. Mice were euthanized seven days post-injury, followed by synovial fluid recovery for global metabolomic profiling and analysis of epiphyseal trabecular bone by micro-computed tomography (μCT). Global metabolomic profiling assessed metabolic differences in the joint response to injury between GF and conventional mice. Magnitude of trabecular bone volume loss measured using μCT assessed early OA progression in GF and conventional mice.

**Results:** μCT found that GF mice had significantly less trabecular bone loss compared to conventional mice, indicating that the GF status was protective against early OA changes in bone structure. Global metabolomic profiling showed that conventional mice had greater variability in their metabolic response to injury, and a more distinct joint metabolome compared to their corresponding controls. Furthermore, differences in the response to injury in GF compared to conventional mice were linked to mouse metabolic pathways that regulate inflammation associated with the innate immune system.

**Conclusions:** These results suggest that the gut microbiota promote the development of PTOA during the acute phase following joint trauma possibly through the regulation of the innate immune system.

## Introduction

Osteoarthritis (OA) is a disease of the whole joint with low-grade articular inflammation playing a role in disease progression[1-4]. Inflammation is associated with age, traumatic injury, and obesity; all of which are OA risk factors[2, 5]. Joint injury triggers an acute inflammatory response immediately after injury, and further perpetuation of inflammation can result in post-traumatic OA (PTOA) via catabolism in the joint[6]. Inflammation post-injury, therefore, is an important contributing factor in OA development, and a better understanding of immune responses in this context may lead to new pharmacological targets for better prevention and treatment.

The perpetual inflammatory response that persists after joint injury includes innate immune activity. Joint injury results in production of endogenous signals termed damage associated molecular patterns (DAMPs) that signal to innate immune cells. In OA, DAMPs include matrix components from cartilage degradation, plasma proteins from vascular leakage, and intracellular alarmins from stressed or necrotic cells[7-10]. Macrophages, neutrophils, and dendritic cells with pattern recognition receptors recognize DAMPs and induce NF-kβ-mediated production of proinflammatory cytokines, such as IL-6, IL-1β, and TNF-α[7-10].

In addition to DAMPs, the immune system is modulated by antigens from microorganisms living in and on the body (microbiome)[11]. Recent studies suggest the gut microbiome may play a role in OA pathogenesis[12, 13] such that innate cells initially activated by joint injury are further and perpetually activated by microbial antigens. For example, increased abundance of *Streptococcus* species in the gut is associated with increased knee pain in OA driven by increased local inflammation in the joint[14]. Moreover, the gut microbiome-produced metabolites, hippurate and trigonelline, distinguish OA progressors from non-progressors[15]. Finally, microbial DNA has been reported in human and mouse cartilage, suggesting that molecules of microbial origin, if not live bacteria themselves, help drive OA-associated inflammation[16]. In a previous study, germ-free (GF) mice (mice lacking a microbiome) had less severe histological OA eight weeks after surgical destabilization of the medial meniscus[17]. However, joint injury induced by surgery requires opening the joint cavity for surgical destabilization, which does not occur in human joint injuries.

Non-invasive PTOA mouse models simulate human injury and avoid some confounding factors from surgical models. Furthermore, non-invasive models allow examination of early timepoints during the acute phase after joint trauma. The acute response to injury is important because it provides a unique opportunity for intervention when patients often seek medical attention. Thus, the objective of this study was to compare the acute responses after joint injury in gnotobiotic and conventional mice. We hypothesize that injured gnotobiotic mice would exhibit less severe OA than conventional mice due to the absence of immune activity driven by the gut microbiome.

## Materials and Methods

### Joint Injury Model and Experimental Design

We utilized a non-invasive injury model to induce PTOA in C57BL/6 mice[18, 19]. This model results in ACL rupture that closely mimics ACL tears in humans and produces a highly reproducible joint injury and associated degeneration of bone and cartilage[18, 19]. Typically, this model first displays changes in subchondral trabecular bone at 7 days with subsequent cartilage pathology detected by 28-56 days post-injury[20].

Mice were anesthetized using isoflurane inhalation and PTOA was induced in conventional and germ-free (GF) mice using a tibial compression system by applying a single compressive overload to the lower leg with a target force of 12N at a loading rate of 130 mm/s[21]. After injury, mice were given a single dose of sustained-release buprenorphine SR (1.0 mg/kg) subcutaneously to alleviate pain associated with ACL rupture. GF mice were maintained in GF-conditions after injury. Naïve uninjured controls without buprenorphine treatment were used for both conventional and GF mice. Mice were euthanized by cervical dislocation seven days post-injury for synovial fluid (SF) collection and whole-knee μCT (Fig. 1).

**Figure 1.**
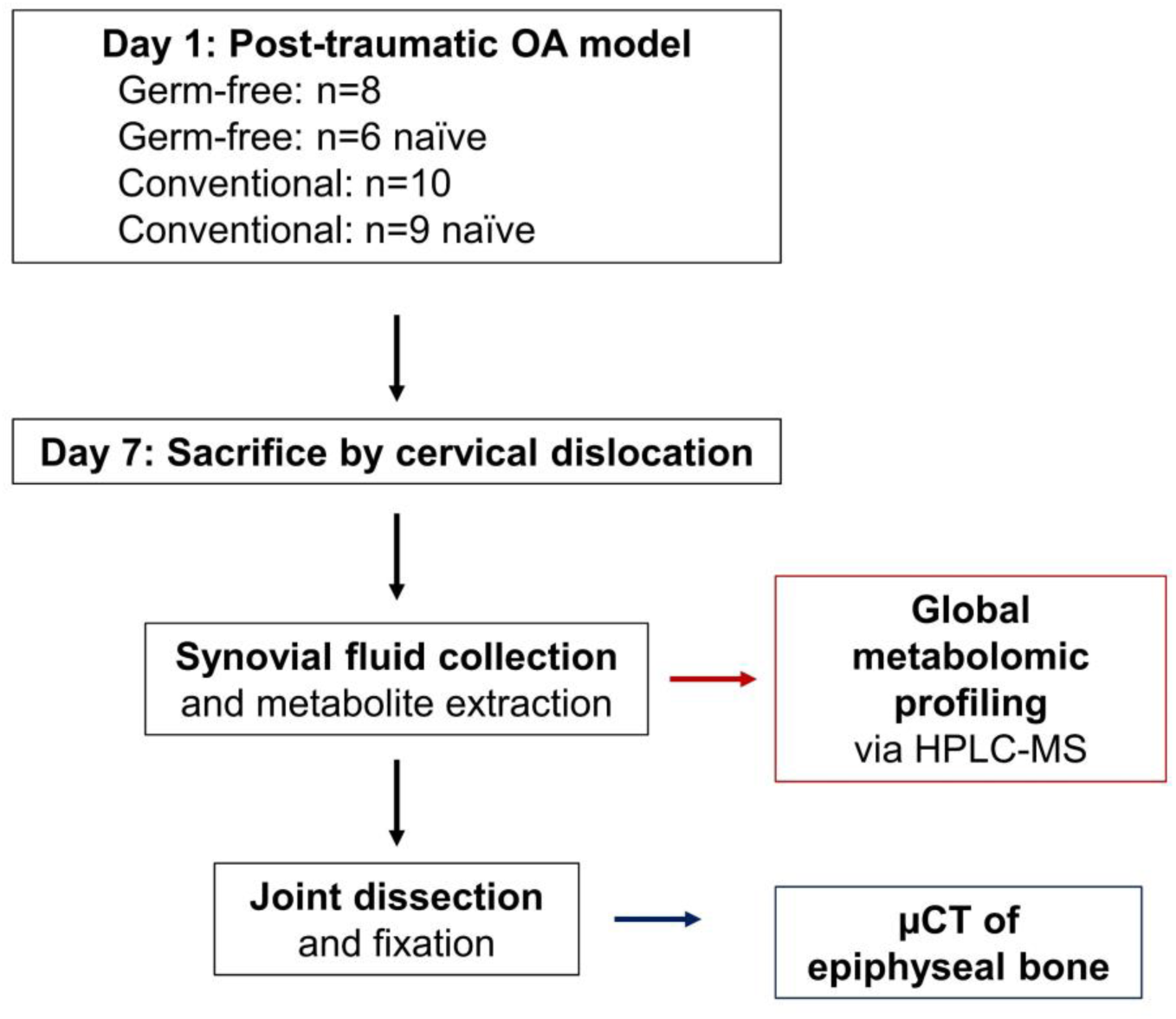
Experimental design to study the role of the microbiome in post-traumatic osteoarthritis. 20-week old C57/BL6 mice were used in this study. Mice were raised in either gnotobiotic (i.e. germ-free) or conventional conditions. On Day 1, mice were subjected to a post-traumatic OA model by single compressive overload of the joint. At Day 7, mice were sacrificed before synovial fluid harvest and joint fixation. Synovial fluid was profiled by global metabolomic analysis, and the epiphyseal bone was analyzed by μCT.

In total, there were six experimental groups for comparisons: GF injured (GF-inj; n=8), GF contralateral (GF-cl; n=8), GF naïve (GF-ctrl; n=5), conventional injured (C57-inj; n=10), conventional contralateral (C57-cl; n=10), and conventional naïve (C57-ctrl; n=9).

Germ-free status was monitored and confirmed using standard cultivation and molecular biology techniques[22]. Briefly, liquid ‘traps’ were established inside hermetically sealed and air purified (HEPA filtered) husbandry isolators containing sterilized drinking water and chow pellets (i.e. the same water and chow that mice were drinking and eating). Traps were monitored daily for any signs of microbial growth. Samples from traps were collected along with stool pellets from mice for aerobic and anaerobic cultivation on non-selective media (Luria broth and Mueller-Hinton broth agar plates) following 48 hours incubation at 37°C. In addition, bulk DNA was isolated from mouse pellets (DNeasy Powersoil Kit, Qiagen, Inc) and attempts were made to amplify a 16S rRNA locus using broad-range (universal) primers[23]. No positive results (*i*.*e*. growth on agar plates or amplification of PCR) were observed.

For injury of GF mice, two cages of animals were removed from the isolator inside a sealed cage with a HEPA-filtered lid and place immediately inside a biosafety cabinet with laminar flow and containing the compressive overload device. All surfaces were contact sterilized ahead of time and the cage was doused repeatedly with disinfectant (Ecolab® Exspor™). Following injury mice were placed back inside the cages and moved back inside husbandry isolators. The isolator environment and mice were checked repeatedly for any indication of breach in sterility as described above.

### Synovial Fluid Collection

Immediately after euthanasia, SF was recovered using the calcium sodium alginate compound (CSAC) method as previously described[24]. Joint cavities were accessed and the patellar tendon was retracted to place a 3mm Melgisorb (Tendra, REF# 250600; Goteborg, Sweden) wound dressing on the cartilage surfaces for SF absorption. The Melgisorb wound dressing was submerged in 35 μL of Alginate Lyase in H_2_O (1 unit/mL concentration; derived from *Flavobacterium*, Sigma-Aldrich A1603-100MG), vortexed, and digested at 34°C for 30 minutes. After digestion, 15 μL of 1.0M sodium citrate (C6H5Na3O7) was added to chelate the Ca^2+^ ions and reduce the viscosity. Sample volumes were recorded prior to freezing at −80°C until metabolite extraction and analysis.

### Metabolite Extraction

Metabolites were extracted from mouse SF following our previously established protocol with minor modifications[25, 26]. SF samples were thawed, centrifuged at 4°C at 500*×g* for 5 minutes, and supernatant was collected for vacuum concentration for ∼2 hr. Metabolites were extracted from the dried pellet by re-suspension in 50:50 water: acetonitrile at −20°C for 30 min. Following extraction, the sample was vortexed for 3 min and centrifuged at 16100*×g* for 5 min at 4°C. Proteins were precipitated by adding 250 μL of acetone and shaken for 3 min followed by overnight refrigeration at 4°C. The mixture was then centrifuged for 5 min at 16100*×g* for 5 min and supernatant was collected for vacuum concentration prior to re-suspension in 50:50 water:acetonitrile for mass spectrometry analysis. All solvents were either HPLC or mass-spectrometry grade.

### Global Metabolomic Profiling

Metabolite extracts were analyzed using established methods[27]. Samples were analyzed on an Agilent 1290 UPLC system (Agilent, Santa Clara, CA, USA) coupled to an Agilent 6538 Q-TOF mass spectrometer (Agilent Santa Clara, CA, USA) in positive mode. The chromatographic run used a Cogent Diamond Hydride HILIC 150 × 2.1 mm column (MicroSolv, Eatontown, NJ) in normal phase with our established elution methods[26]. Spectra were analyzed as previously described[26].

### μCT Analysis of Subchondral Bone

Following dissection, whole knee joints were fixed in 4% paraformaldehyde for 24-48 hours and preserved in 70% ethanol. Joints were individually imaged using micro-computed tomography (SkyScan 1173, Bruker, Belgium) to quantify trabecular bone microstructure of the distal femoral epiphysis. During imaging, bones were embedded in 1.5% agarose, and were scanned with a 8.1 μm nominal voxel size (x-ray tube potential = 40 kVp, current = 200 μA, integration time = 1300 ms) according to the JBMR guidelines for μCT analysis of rodent bone structure[28]. Transverse images were exported as bitmap image files, then converted to raw files using ImageJ and uploaded to another μCT system for analysis (SCANCO μCT 35, Brüttisellen, Switzerland). The trabecular bone volume of interest was contoured by manually drawing on transverse images, and included the full epiphyseal region, excluding cortical bone and the growth plate (Fig. 2). Trabecular bone volume fraction (BV/TV), trabecular thickness (Tb.Th), and other microstructural outcomes were determined using the SCANCO analysis software.

**Figure 2.**
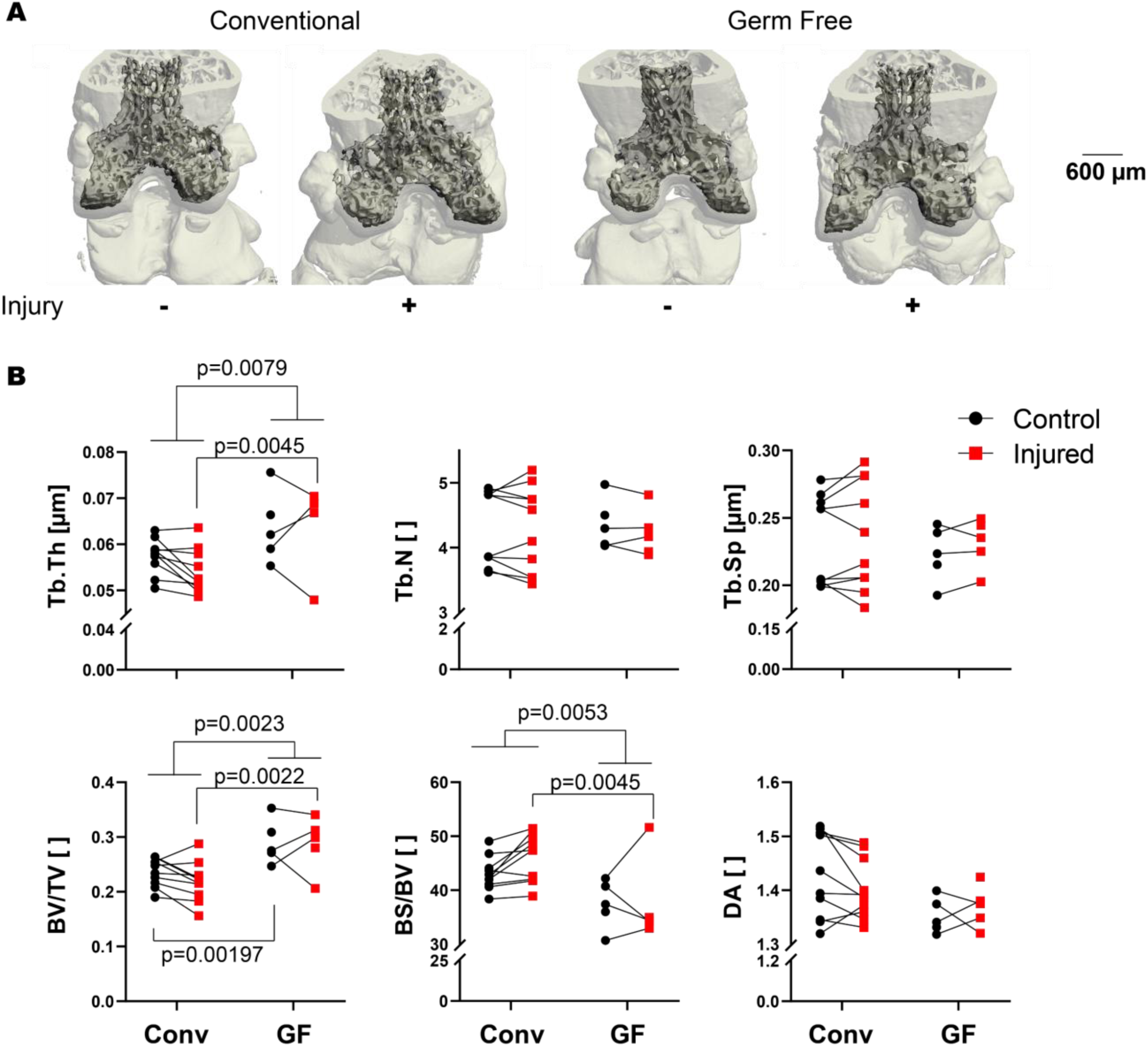
Germ-free mice have increased epiphyseal bone mass compared with conventional controls in both injured and contralateral knees. (A) Representative μCT reconstructions of conventional and germ-free femurs. (B) Microstructural parameters of femoral epiphysis quantified by μCT. Germ free mice had greater trabecular thickness (Tb.Th, p=0.0079), and injury induced thicker trabecular in GF mice (p=0.045). Germ free mice also had greater volume fraction than conventional mice independent of injury, as well as in both control and injured knees (all p<0.0022). Numbers of trabeculae, trabecular spacing, and degree of anisotropy (DA) were similar between GF and conventional mice.

### Statistical Analyses

Before statistical analyses, median metabolite intensities were calculated for each metabolite across each experimental group. Metabolite features (m/z values) with a median intensity of zero across all experimental groups were removed from further analyses. Any remaining metabolites with intensity of zero (*i*.*e*. non-detected) were replaced with one-half the minimum peak intensity for statistical analyses.

Statistical analyses were performed in MetaboAnalyst and GraphPad Prism (GraphPad Prism Software, La Jolla, CA, USA) for metabolomics data and μCT data, respectively[29]. Metabolomics data were log transformed (log2) and standardized (mean centered divided by standard deviation) in MetaboAnalyst prior to statistical analyses[29]. All statistical tests used a false discovery rate (FDR)-corrected *a priori* significance level of 0.05.

The metabolomic profiles were analyzed using multivariate statistical methods to determine between-group changes and patterns of co-regulated metabolites. Unsupervised statistical methods, hierarchical cluster analysis (HCA) with Euclidean distances, and principal component analysis (PCA) were used to examine variation within the overall dataset. PCA was visualized with scatterplot projections onto the principal axes to examine sample to sample variation.

Differentially regulated metabolites were identified using two-tailed Student’s t-tests, analysis of variance (ANOVA) F-tests, and volcano plot analysis. While Student’s t-tests and ANOVA identify metabolites that were significantly different by FDR-corrected p-values, volcano plot assesses significance and magnitude of change simultaneously (FDR-corrected p-value<0.05 and fold-change greater than twofold).

Specific metabolite features of interest were matched to putative metabolite identities using the MetaboAnalyst “MS Peaks to Pathways” function[29, 30]. This generates a compound match list, which matches *m/z* values to possible metabolite identities, while simultaneously determining enriched metabolic pathways within the relevant group of metabolite features.

## Results

### Less evidence of OA-related injury-induced bone loss in germ-free mice

We found no evidence that GF mice became colonized by any microorganism up to eight days following injury. μCT analysis showed that GF mice had greater trabecular bone volume than conventional mice, with 23% greater trabecular bone volume fraction (BV/TV) and 11% greater trabecular thickness (Tb.Th, Figure 4). GF mice had thicker trabeculae independent of injury (overall ANOVA, p=0.0079) and injured GF mice had thicker trabeculae than conventional mice (p=0.0045). Bone volume fraction was also greater in GF mice than conventional independent of injury status (overall ANOVA, p=0.0079). Both the contralateral and injured knees of GF mice had greater BV/TV than conventional mice (p=0.00197 and p=0.0022, respectively). There were no differences in trabecular number (TB.N) or trabecular separation (Tb.Sp) between GF and conventional mice.

**Figure 3.**
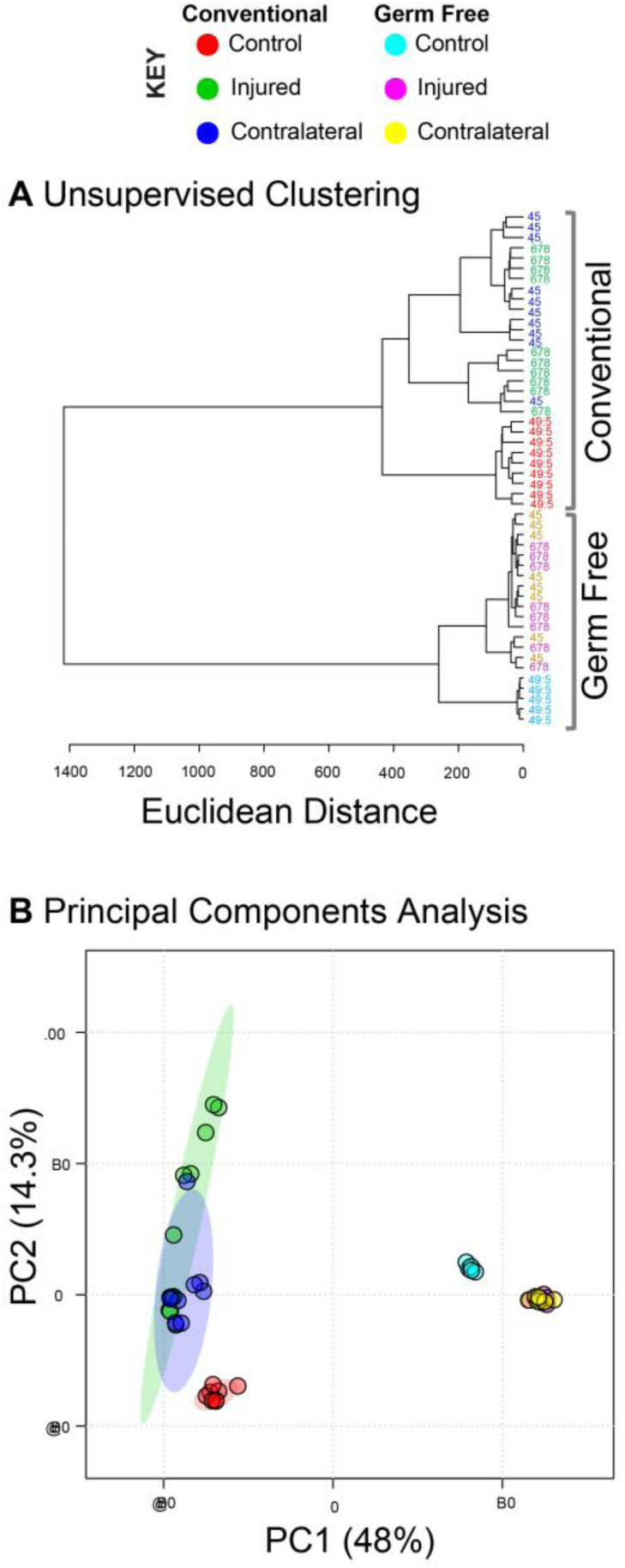
Global metabolomic profiles are significantly different between germ-free and conventional mice. The total of 13,488 metabolites were subjected to both unsupervised hierarchical clustering and principal components analysis. (A) Unsupervised hierarchical clustering finds distinct clusters between the metabolomic profiles of germ-free and conventional mouse synovial fluid. Furthermore, within both groups the naïve controls were distinct from both the contralateral and injured knees of injured mice. (B) Principal components analysis indicates that both injury and germ-free status drive differences in global metabolomic profiles.

**Figure 4.**
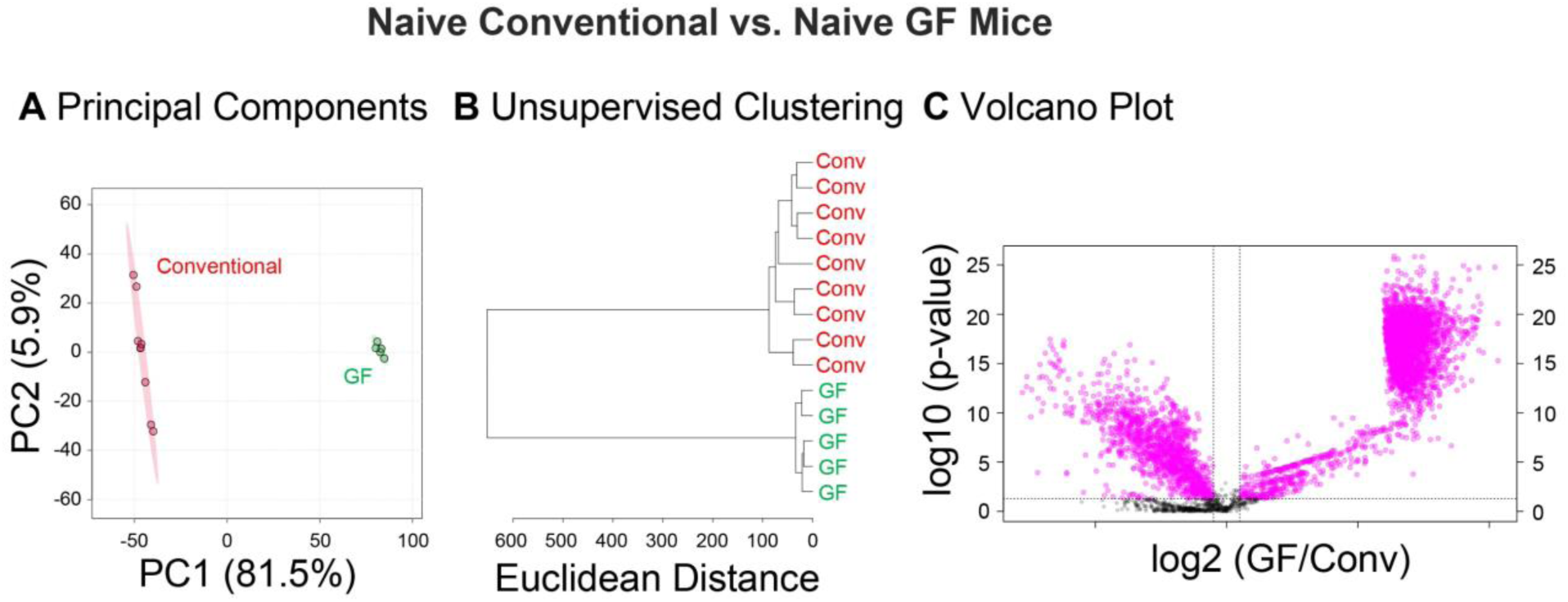
Global metabolomic profiles differ between naïve knees of germ free and conventional mice. (A) Principal components analysis finds clear separation between synovial fluid metabolomic profiles of germ-free and conventional mice. (B) Unsupervised clustering finds distinct clusters between conventional and germ-free mice. (C) Volcano plot analysis found 4357 metabolites significantly different between germ-free and conventional mice.

On average, injured joints of conventional mice had a 7.4% lower BV/TV compared to their contralateral limbs, although this difference was not statistically significant (p = 0.061). Injured joints of conventional mice also had 5.9% lower Tb.Th (p = 0.016) and 7.0% greater BS/BV (p = 0.015) than contralateral joints. In contrast, no significant differences in trabecular bone microstructure were observed between injured and contralateral joints from GF mice.

### Germ-free mice had distinct metabolomic profiles compared to conventional mice, injured and uninjured

Global metabolomic profiling of mouse SF detected 13,488 metabolite features across all experimental groups. 4,689 metabolite features were identified as significant by ANOVA (FDR-adjusted p-values<0.05). Unsupervised analyses including PCA and HCA found metabolomic variation between mouse SF cohorts, while FDR-corrected t-tests, and volcano plot analyses identified differentially regulated metabolites.

PCA and HCA revealed two large clusters that corresponded to GF and conventional mice, confirming that the presence of the gut microbiome influences the overall joint metabolome (Fig. 3). Conventional mice displayed greater overall variability as shown by increased distance between samples in PCA ordination space and increased branch lengths in HCA dendrogram (Fig. 3). Similarly, comparing synovial fluid of naïve, uninjured mice (Fig. 4), there was dramatically less variability in the GF mice. Unsupervised clustering resulted in complete separation of GF and conventional samples (Fig. 4B). Taken together, these results showed that the presence of a gut microbiome produces a distinct joint metabolome with greater variability between individuals.

After injury, the metabolomes of both GF and conventional mice shifted markedly from their naïve, uninjured counterparts (Fig. 3). Furthermore, there was significant overlap between injured and contralateral samples in both conventional and GF mice, suggesting injury produces a systemic response that influences both joint metabolomes.

Metabolomic profiles also suggested that GF mice were less sensitive to injury than conventional mice. For example, branch lengths between dendrogram clusters represent differences in global metabolomic profiles, or dissimilarity. Within conventional mice, naïve individuals had a dissimilarity (Euclidean distance) of approximately 450 from injured mice (Fig. 3A). The same comparison in GF mice yielded a dissimilarity of only 275, suggesting that the lack of a microbiome results in decreased metabolomic plasticity following injury.

### Joint metabolome in germ-free and conventional naïve mice

To confirm that GF mice have a distinct joint metabolome than conventional mice in uninjured control conditions, a pairwise analysis of naïve experimental groups was conducted (GF-ctrl vs. C57 ctrl). A total of 12,306 metabolite features were detected between GF and conventional naïve cohorts, with 4382 features identified as significantly different by Student’s T-test (FDR-adjusted p-value<0.05).

Global differences between GF and conventional naïve mice were assessed with unsupervised PCA and HCA. Unsupervised analyses showed clustering of SF samples within their respective cohorts, with greater variability within the C57-ctrl cohort than GF-ctrl (Fig. 4A-B). Specific differences were identified using volcano plot analysis. Volcano plot analysis identified 3078 metabolite features as significantly upregulated and 1279 metabolite features as significantly downregulated in GF-ctrl compared to C57-ctrl mice (Fig. 4C).

### Metabolic response to injury in germ-free injured and conventional mice

To assess the distinct response to injury in GF compared to conventional mice, pairwise analyses were conducted using only injured experimental groups (GF-inj vs. C57-inj). A total of 12,075 metabolite features were detected between GF and conventional injured cohorts, with 3552 features identified as significantly different by Student’s T-test (FDR-adjusted p-value<0.05).

Global differences between GF and conventional injured mice were assessed with unsupervised PCA and HCA. Both PCA and HCA showed distinct clustering of SF samples within their respective cohorts (Fig. 5A-B). However, PCA and HCA once again showed greater variability within the C57-inj mice cohort as shown by less tight clustering of samples compared to GF-inj samples (Fig. 5A-B).

**Figure 5.**
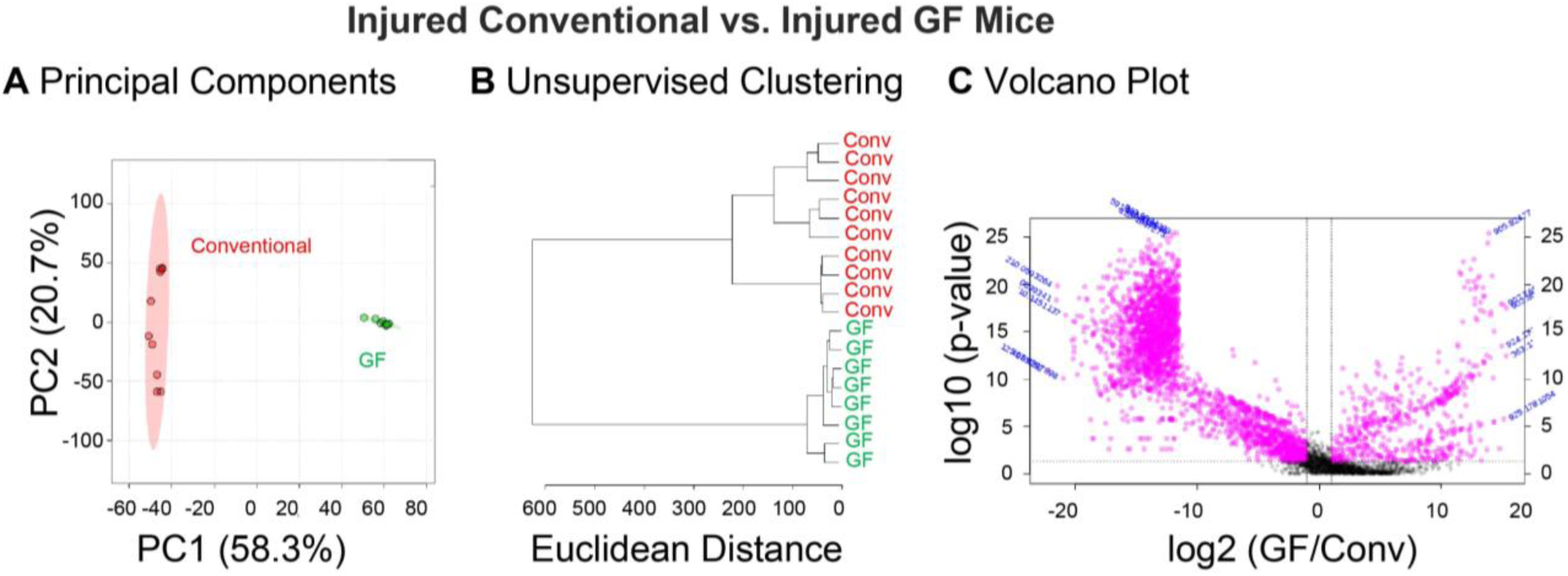
Global metabolomic profiles of synovial fluid are driven by gut microbiota. Synovial fluid metabolites compared between naïve germ free and conventional mice. (A) Principal components analysis finds complete separation between germ-free and conventional metabolomic profiles. (B) Germ free and conventional mice separate into two distinct clusters by unsupervised hierarchical clustering analysis. (C) Volcano plot analysis shows substantial differences in individual metabolites between the synovial fluid of germ free and conventional mice.

Volcano plot analysis was used to identify specific metabolite features that were significantly upregulated or downregulated in GF compared to conventional injured mice. 2728 metabolite features were significantly downregulated, and 578 metabolite features were significantly upregulated in GF-inj compared to C57-inj mice (Fig. 5C). Metabolite features that were significantly downregulated in GF-inj compared to C57-inj mice mapped to arachidonic acid metabolism, the TCA cycle, porphyrin metabolism, arginine and proline metabolism, and pyruvate metabolism (Table 1). Metabolite features significantly upregulated in GF-inj compared to C57-inj mapped to valine, leucine, and isoleucine degradation, porphyrin metabolism, glycerophospholipid metabolism, arachidonic acid metabolism, fatty acid degradation, N-glycan biosynthesis, propanoate metabolism, and pantothenate and Coenzyme A (CoA) biosynthesis (Table 1).

**Table 1.** Pathways altered in Germ-Free mice. Metabolite features identified by volcano plot analysis as significantly upregulated or downregulated in Germ Free mice compared with conventional were mapped to KEGG metabolic pathways. Pathways are reported with directionality (upregulated or downregulated); the total number of metabolites in the pathway, total number of metabolite features detected in samples in that pathway, total number of significant metabolite features (significant by both FDR-corrected p-value and fold change), and p-value for enrichment. Most significant pathways are reported up until arbitrary p-value cut-off for upregulated (p-value < 0.05) and downregulated pathways (p-value < 0.1).

| Pathway | Up/Down | Total<br>metabolites<br>in pathway | Total<br>pathway<br>hits | Significant<br>pathway<br>hits | P-value |
| --- | --- | --- | --- | --- | --- |
| Valine, leucine and isoleucine<br>degradation | ↑ | 40 | 16 | 8 | 0.012306 |
| Porphyrim metabolism | ↑ | 30 | 12 | 5 | 0.020232 |
| Glycerophospholipid<br>metabolism | ↑ | 36 | 9 | 4 | 0.022792 |
| Arachidonic acid metabolism | ↑ | 36 | 13 | 5 | 0.023064 |
| Fatty acid degradation | ↑ | 39 | 7 | 3 | 0.033277 |
| N-Glycan biosynthesis | ↑ | 41 | 3 | 2 | 0.036671 |
| Propanoate metabolism | ↑ | 23 | 8 | 3 | 0.040544 |
| Pantothenate and CoA<br>biosynthesis | ↑ | 19 | 8 | 3 | 0.040544 |
| Arachidonic acid metabolism | ↓ | 36 | 13 | 13 | 0.062431 |
| Citrate cycle (TCA cycle) | ↓ | 20 | 12 | 12 | 0.07051 |
| Porphyrim metabolism | ↓ | 30 | 12 | 12 | 0.07051 |
| Arginine and proline metabolism | ↓ | 38 | 10 | 10 | 0.091788 |
| Pyruvate metabolism | ↓ | 22 | 10 | 10 | 0.091788 |

## Discussion

The goal of this study was to establish new GF mouse modeling techniques that can be used to experimentally evaluate host response to acute OA-associated injury in the absence of a microbiome. Since we found no evidence that GF mice became colonized by microorganisms following injury, this study provides direct proof-of-principle that such experimental manipulations can be achieved in a sterile setting. We feel confident that similar approaches can be used to further examine gnotobiotic, or defined microbiome, impacts on host response following joint injury.

Inflammation clearly plays a role in the development of OA in humans[1-4] as well as in the development of OA risk factors, including age, obesity, and joint trauma. After joint trauma there is an initial acute inflammatory response followed by low-grade chronic inflammation[6]. Initial inflammatory responses from DAMPs promote a state of stress[31, 32], and it is becoming increasingly clear that other endogenous signals from the microbiome further exacerbate pro-inflammatory responses[11, 33].

Bacteria in the mammalian gut play key roles in many diseases associated with chronic inflammation, including type 2 diabetes, inflammatory bowel disease, rheumatoid arthritis, and ankylosing spondylitis[34-37]. Given that chronic inflammation is now an established contributor to OA progression, a better understanding of the impact of the overall gut microbiome and that of individual members as activators (pro-inflammatory) and regulators (anti-inflammatory) of OA-related inflammation promises to yield new targets for prevention and treatment. Furthermore, because joint trauma increases OA risk and because these patients typically only seek care shortly after injury, determining how the gut microbiome influences PTOA over a longer recovery period is critically important.

It is well-known that immunity in GF mice is markedly different from conventional counterparts with observed deficits in overall numbers and distribution of leukocytes[38]. These deficits lead to altered acute responses to stress in other injury-induced models[39], including lymph node trafficking. Similarly, increased bone loss is associated with inflammatory diseases via an imbalance in bone resorption and formation [40].

In our study, synovial fluid metabolomes between injured and naïve mice were more distinct from one another in conventional compared to GF mice. This was further supported by μCT data. Collectively these data suggest that the GF phenotype is protective against the development of OA and the lack of a microbiome may slow the progression of disease. Metabolomic analyses also showed greater variability within conventional versus GF mice, which is consistent with known differences in immune cell activation/maturation and in turn, responses to a diverse community of microorganisms.

Moreover, the observed alterations in joint metabolomic profiles are consistent with different inflammatory responses in conventional versus GF mice. For example, fatty acid degradation, pantothenate and CoA biosynthesis, and arachidonic acid metabolism identified in volcano plots are known to play key roles in inflammation[41, 42]. Cellular fatty acid degradation is typically accomplished by beta oxidation, which is an important anti-inflammatory mediator of innate immunity[43]. Interestingly, this pathway was significantly upregulated in GF versus conventional mouse joint metabolome. While these results suggest that the absence of a microbiome promotes anti-inflammatory processes via fatty acid degradation, other studies report that short-chain fatty acids (SCFAs) produced by the microbiome also suppress immune functions[44]. Therefore, the effect of SCFAs may not be sufficient to counterbalance the skewed immune responses of GF mice.

Pantothenate, also known as vitamin B5, and arachindonic acid (AA) were other known mediators of the innate immunity associated with the GF joint metabolome. One study found that B5 had pro-inflammatory effects during *Myobacterium tuberculosis* infection[45], while another study found the overall vitamin B complex (B1, B2, B3, B5, B6, B12) suppressed local inflammation after peripheral nerve transection[46]. Furthermore, B5 is a precursor to CoA, which plays a role in regulating the innate immune system[47]. AA is released from the lipid bilayer in response to injury or stress and initiates a series of signaling cascades that can trigger inflammation[48]. While our results suggest these pathways help explain observed differences in response to joint injury in the context of a microbiome, neither have received much attention with respect to OA development.

An important difference between previously published approaches and ours is the use of the non-invasive PTOA mouse model, which has been well-established to initiate rapid loss of trabecular bone as early as day seven post-injury[20]. Microstructural differences between injured and contralateral joints observed in conventional mice were not observed in GF mice, supporting the conclusion that immune deficits in mice lacking a microbiome protected them against rapid trabecular bone loss that is typical in this PTOA mouse model.

This work has important limitations. First, GF mice are raised in isolators, which can alter behavior[49]. Thus, differences observed between conventional and GF mice may result from additional factors besides the lack of a microbiome, such as differences in activity levels or body composition that could potentially confound our results. Second, while the CSAC method of rodent synovial fluid collection is established[24], this method can be challenging. In preliminary studies, this method consistently yielded thousands of metabolites compared with blank and neat controls indicating that synovial fluid metabolites can pe profiled using the CSAC method. Third, per IACUC requirements, injured mice received pain medication after injury. While pharmacokinetics studies[50] suggest that plasma concentrations of sustained-release buprenorphine diminish before the 7-day endpoint, the observed effects of the GF response to injury might have been greater without buprenorphine. Fourth, while all mice limped after injury, ACL tears were not confirmed through laxity testing.

In conclusion, to our knowledge this study is the first to investigate the role of the microbiome in the acute phase after joint injury. These results demonstrate that the gut microbiome drives key injury responses, specifically via perturbations in the innate inflammatory response to injury. Thus, microbiota may be a potential therapeutic target during the acute phase after joint trauma to abrogate the innate immune response and thereby diminish some of the first events that result in PTOA.

## Acknowledgements

We thank Dr. Tim Griffin for showing us the CSAC method, and we acknowledge Dr. Ladean McKittrick for his assistance with μCT imaging. We thank the Montana State University Proteomics, Metabolomics, and Mass Spectrometry Facility for assistance with sample analysis. This facility is supported in part by funding from the Murdock Charitable Trust and NIH P20GM103474 of the IDEA program and AG007996. This research is funded by the NSF (CMMI 1554708 to RKJ), NIH (R01AR073964 to RKJ, R01AR075013 to BAC), Murdock Charitable Trust (FSU-2017207 to AKH), and Montana Space Grant Consortium for student funding.

## Author contributions

AKH performed the injury model, extracted metabolites, analyzed data, and drafted the manuscript. CWW assisted in the injury model and joint dissections. HDW analyzed data. EB assisted with joint dissections and metabolite extractions. MM cared for GF mice and assisted with the injury model. BAC analyzed and interpreted μCT data. STW assisted with GF mouse care, designed experiments, and analyzed data. RKJ designed experiments and analyzed data. All authors have read and revised the manuscript.

## Figures and Tables

F1. Exp design

F2. MicroCT

F3. Global metabolomic Profiles

F4. Control Conventional vs Control GF

F5. Injured C57 vs Injured GF

T1. Pathways

## Supplemental Tables

ST1. Mouse code/ids

ST2. All metabolite data

ST3. Pathways (current ST2)

ST4. Pathways (current ST3)

## Supplemental Material

**Supplemental Table 1. Mouse information from experimental cohorts**. Mouse labels, type (germ-free or C57BL/6), sex (male or female), weight (grams), and leg status (injured, contralateral, naïve).

**Supplemental Table 2. Metabolomics data**. Original metabolomics data generated by HPLC-MS formatted for MetaboAnalyst. Metabolite features with a median value of zero across all groups were removed.

**Supplemental Table 3. Full enrichment analysis of significantly upregulated metabolite features identified by volcano plot analysis**. Pathways are reported with the total number of metabolites in the pathway, total number of metabolite features detected in samples in that pathway, total number of significant metabolite features (significant by both FDR-corrected p-value and fold change), expected number of metabolites in the pathway, Fisher’s exact p-value for the pathway (FET), EASE score (modified Fisher’s exact p-value), and Gamma-adjusted p-value for the pathway.

**Supplemental Table 4. Enrichment analysis of significantly downregulated metabolite features identified by volcano plot analysis**. Pathways are reported with the total number of metabolites in the pathway, total number of metabolite features detected in samples in that pathway, total number of significant metabolite features (significant by both FDR-corrected p-value and fold change), expected number of metabolites in the pathway, Fisher’s exact p-value for the pathway (FET), EASE score (modified Fisher’s exact p-value), and Gamma-adjusted p-value for the pathway.

